# Mosaic evolution of molecular pathways for sex pheromone communication in a butterfly

**DOI:** 10.1101/2020.09.04.279224

**Authors:** Caroline M. Nieberding, Patrícia Beldade, Véronique Baumlé, Gilles San Martin, Alok Arun, Georges Lognay, Nicolas Montagné, Lucie Bastin-Héline, Emmanuelle Jacquin-Joly, Céline Noirot, Christophe Klopp, Bertanne Visser

## Abstract

Unraveling the origin of molecular pathways underlying the evolution of adaptive traits is essential for understanding how new lineages emerge, including the relative contribution of conserved, ancestral traits, and newly evolved, derived traits. Here, we investigated the evolutionary divergence of sex pheromone communication from moths (mostly nocturnal) to butterflies (mostly diurnal) that occurred ~98 million years ago. In moths, females typically emit pheromones to attract male mates, but in butterflies pheromones and used by females for mate choice. The molecular bases of sex pheromone communication are well understood in moths, but have remained virtually unexplored in butterflies. We used a combination of transcriptomics, real time qPCR, and phylogenetics, to identify genes involved in different steps of sex pheromone communication in the butterfly *Bicyclus anynana*. Our results show that the biosynthesis and reception of sex pheromones relies both on moth-specific gene families (reductases) and on more ancestral insect gene families (desaturases, olfactory receptors, odorant binding proteins). Interestingly, *B. anynana* further appears to use what was believed to be the moth-specific neuropeptide Pheromone Biosynthesis Activating Neuropeptide (PBAN) for regulation of sex pheromone production. Altogether, our results suggest that a mosaic pattern best explains how sex pheromone communication evolved in butterflies, with some molecular components derived from moths, and others conserved from more ancient insect ancestors. This is the first large-scale analysis of the genetic pathways underlying sex pheromone communication in a butterfly.

## Introduction

Evolution of new life forms occurs through the transition from an ancestral to a descendant clade, where the newly formed clade generally expresses a mosaic of traits conserved from the ancestor, as well as newly evolved, derived, traits. Mosaic evolution is indeed a recurring pattern in paleontology (Cracraft 1970; Gómez-Robles *et al.* 2014; Xu *et al.* 2017). For example, *Tiktaalik roseae*, believed to represent the transition from fishes to amphibians (the “fishapod”; ~375 Mya), shares some traits with more primitive sarcopterygian fishes (e.g. body scales, fin rays, lower jaw and palate) and other traits more typical of tetrapods (e.g. shortened skull roof, a modified ear region, a mobile neck, a functional wrist joint, and other features) (Daeschler *et al.* 2006). Investigating the genetic bases of ancestral and derived phenotypic traits is essential to obtain a mechanistic explanation of how mosaic evolution takes place. Examples of studies investigating the mechanistic basis of mosaic evolution have accumulated in the last decade, including recent genomic evolution analyses identifying patterns of gene loss, retention, and *de novo* evolution (e.g. van Gestel *et al.* 2019; Fernández & Gabaldón 2020; Guijarro-Clarke *et al.* 2020). Other patterns in the genetic bases of trait evolution have suggested a role for hybridization between species (e.g. Berner & Salzburger 2015; Stryjewski & Sorenson 2017; Marques *et al.* 2019) or co-option of molecular pathways that acquired new functions (e.g. Shubin et al. 2009, including a butterfly, Shirai et al. 2012). Derived phenotypic traits can thus be generated through different molecular mechanisms that need to be identified case-by-case.

Here, we focused on the genetic bases of divergence in sex pheromone communication during the evolutionary transition from moths to butterflies, which occurred ~98 Mya. Sex pheromone communication is used by most insects, including butterfly species, for finding, identifying, and assessing the quality of potential mating partners (Myers 1972; Birch *et al.* 1990; Vane-Wright & Boppre 1993; Nieberding *et al.* 2008; Sarto Monteys *et al.* 2016). Sex pheromone communication is under strong selection, because it determines mating success and consequently an individual’s contribution to the next generation (Svensson 1996; Moore *et al.* 1997; Smadja & Butlin 2009; Hansson & Stensmyr 2011; Wyatt 2014). Molecular pathways for sex pheromone biosynthesis, its regulation, and pheromone reception have been identified in several moth species (Tillman *et al.* 1999; Jurenka 2004; Blomquist *et al.* 2012; Leal 2013; Yew & Chung 2015; Zhang *et al.* 2015; Rafaeli 2016). Compared to other insects, moths appear to have evolved moth-specific genes or gene lineages involved in sex pheromone communication (Jurenka 2004; Pelosi *et al.* 2006; Vieira *et al.* 2007; Leal 2013; de Fouchier *et al.* 2014; Helmkampf *et al.* 2015; Yew & Chung 2015; Bastin-Héline *et al.* 2019). For biosynthesis, most moths use a limited number of enzymes for desaturation, chain-shortening, reduction, acetylation or oxidation of *de novo* synthesized saturated fatty acids (Jurenka 2004; Liénard *et al.* 2014; Helmkampf *et al.* 2015; Yew & Chung 2015), which generate tremendous chemical diversity of pheromone components and high species-specificity of pheromone blends (Tillman *et al.* 1999; Jurenka 2004). Some of these enzyme families comprise Lepidoptera-specific subclades, such as the Δ9- and Δ11-desaturases (Jurenka 2004; Yew & Chung 2015; Helmkampf et al 2014). Similarly, novel and derived odorant receptor and odorant binding protein subclades have evolved in moths, which bind specifically to sex pheromone chemicals (Pelosi *et al.* 2006; Vieira *et al.* 2007; Leal 2013; de Fouchier *et al.* 2014; Bastin-Héline et al 2019). We aimed to investigate whether butterflies use moth-specific molecular pathways, more ancestral insect pathways, or have evolved butterfly-specific molecular pathways for pheromone communication.

Butterfly sex pheromones have some ecological specificities that contrast with those of moths, for which it is used by female moths to signal their location to mating partners in the dark or at dusk. In contrast, butterfly sex pheromones are generally produced by males and the importance of sex pheromone communication in butterflies was less clear due to their diurnal lifestyle. Studies on some butterfly species have, however, revealed that sex pheromone communication is important for determining mating success (e.g. Costanzo and Monteiro 2007; Nieberding et al. 2008) and fuels speciation events (Bacquet *et al.* 2015). Butterfly sex pheromones can convey refined information with regard to the identity or quality of potential mating partners and can be critical for mate choice and species recognition (Andersson 1994; Johansson & Jones 2007). We focused on the butterfly *Bicyclus anynana*, whose sex pheromone composition has been previously identified and functionally validated (Nieberding *et al.* 2008, 2012). Moreover, experimental manipulation, including the addition of synthetic sex pheromone perfumes (Nieberding *et al.* 2008, 2012), and artificial induction of “anosmia” (Costanzo & Monteiro 2007; van Bergen *et al.* 2013), confirmed the importance of *B. anynana* sex pheromone for mating success. More than a hundred chemical components have been identified on *B. anynana* adult male and female bodies (Heuskin *et al.* 2014), but the composition of the male sex pheromone (“MSP” hereafter) consists of three main volatile components: (Z)-9-tetradecenol (MSP1), hexadecanal (MSP2) and 6,10,14-trimethylpentadecan-2-ol (MSP3) (Nieberding *et al.* 2008). The identification and functional validation of these three MSPs and their role in *B. anynana* chemical communication set this species apart from other lepidopterans, where the role of specific chemical components often remains elusive in relation to fitness (but see Nieberding PLoS ONE 2008 where specific components were identified). MSP1 and MSP2 are derived from fatty acids and are typically found in the pheromone blends of many moth species (Jurenka 2004), which led us to hypothesize that *B. anynana* uses the same genes as moths for sex pheromone communication.

To investigate if butterflies use what were believed to be moth-specific gene families, despite 98 million years since the divergence of butterflies from moths (Wahlberg *et al.* 2013), we used RNA-seq to identify genes that could be involved in *B. anynana* pheromone production, biosynthesis regulation, and reception. We compared transcript abundance of sex pheromone-related adult tissues (male pheromone producing structures, heads and antennae) with control tissues (female wings and heads), and validated our findings with qPCR. We identified specific candidate genes involved in the different olfactory communication functions and used phylogenetic analyses to identify the molecular origin of those genes. Our results reveal that sex pheromone communication in *B. anynana* evolved through a mosaic of ancestral insect genes, and more derived lepidoptera-specific genes.

## Results and Discussion

### Candidate gene identification by transcriptome sequencing

Numerous publications document gene expression studies focusing on chemical communication in Lepidoptera, but only two studied butterflies (Briscoe *et al.* 2013; van Schooten et al 2016), and none of those investigated butterfly sex pheromone communication. Here, we produced six RNA libraries from different adult tissues that were specifically chosen to cover the various steps of male pheromone communication (Figure 1): pheromone biosynthesis (which occurs in dedicated structures on male wings, called androconia; Nieberding *et al.* 2008), its neuro-regulation (in the brain), and pheromone reception (in antennae). Approximately 500 male and female *B. anynana* adults were dissected and relevant tissues assigned to different libraries (Figure 1A). For pheromone synthesis, we compared transcripts in male androconia (Library “androconia”) with those in remaining adult male wing parts (Library “male wings”) and adult female wings (Library “female wings”) as controls. For regulation of pheromone communication, we compared transcript abundance between adult male heads (where the regulation of pheromone synthesis takes place; Library “adult male heads”) and adult female heads (Library “adult female heads”, control). For pheromone reception, we compared transcripts between adult male and female antennae (the tissue where pheromone reception takes place; Nieberding et al (2008); Library “antennae”) and adult heads (Libraries “male heads” and “female heads”) as controls. Two other libraries were also analyzed, corresponding to pupal wings in males (Library “pupal male wings”), and females (Library “pupal female wings”), but these data will not be discussed to focus solely on adults, the stage during which pheromone communication takes place. The total of 737,206 Roche 454 reads obtained from the different tissues sampled in *B. anynana* were assembled into 43,149 contigs, with 76,818 remaining non-assembled singletons (Figure 1B and C, Supplementary Table 1). The transcriptome contained 48.6% near-complete sequence information compared to the insect database (S: 43%; D: 5.6%; F:23%; M: 28.4%; BUSCO v4.0.6; Simão *et al.* 2015). The transcripts were annotated based on reference genomes for several butterfly species (including *B. anynana*; Nowell *et al.* 2017), as well as other relevant insect species (see materials and methods section ‘Transcriptome assembly, quantification, and annotation’).

**Figure 1:**
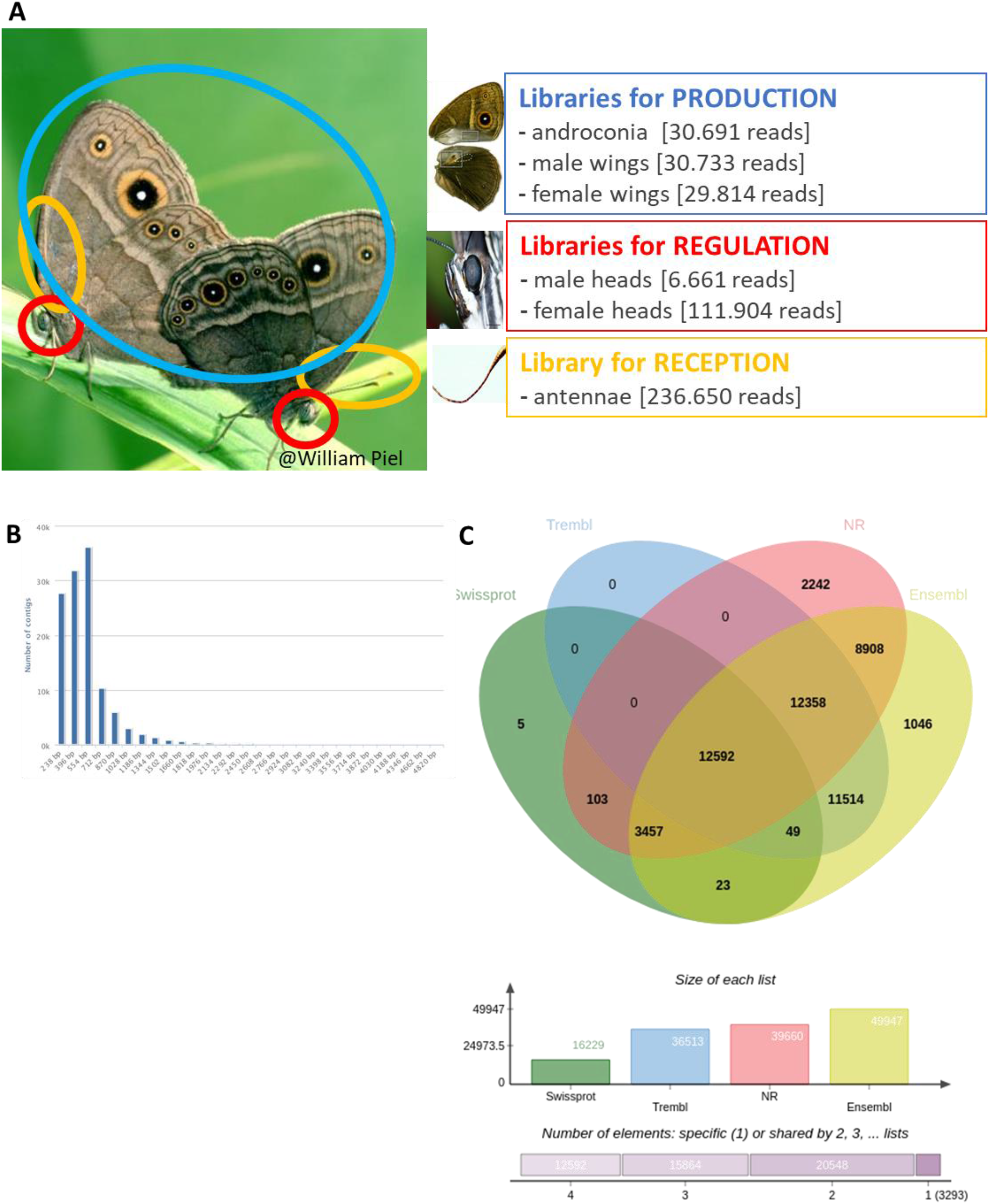
A) Experimental design for the transcriptomics experiment showing the 3 steps involved in male sex pheromone communication and the corresponding tissues sampled to produce the RNA libraries (also including developmental libraries, not shown here). The number of sequenced and cleaned reads per library is provided. 8) Information about the number of contigs in the transcriptome. C) Venn diagram of annotated contig with regard to databases: Swissprot, Trembl, NR and 10 species of Ensembl Lepbase.

Using the digital differential display (DDD) tool (of NCBI’s UniGene database; p < 0.05), a total of 422 contigs were found to be differentially expressed when tissue-specific libraries were compared (Supplementary Table 1). Expression differences were validated by real time quantitative PCR analyses on 10 selected candidate chemical communication genes, showing that relative differences in expression levels in our transcriptome dataset matched those quantified by RT-qPCR (Supplementary Figure 1).

### B. anynana *sex pheromone biosynthesis*

*B. anynana* was the first butterfly for which sex pheromone biosynthesis molecular pathways were investigated and compared to those of moths (Liénard *et al.* 2014). One gene related to pheromone communication was overexpressed in *B. anynana* male androconial wing tissues compared to male and female control wing samples (i.e., ‘type 1’ contigs in Supplementary Table 1), an aldose reductase-like gene. This gene was also overexpressed in the male androconial wing tissue alone. Moreover, a Δ9-desaturase gene was also found to be overexpressed in this library. In contrast to earlier findings in *B. anynana* (Liénard et al. 2014), no fatty-acyl reductase (FAR), nor Δ11-desaturase were found overexpressed in male androconial wing tissue (Supplementary Table 1).

Desaturases (that add a double bond to fatty acid substrates) were previously found to be involved in *B. anynana* MSP biosynthesis (Liénard *et al.* 2014). We, therefore, extended our search for desaturase genes for each of the libraries separately. We focused specifically on Δ9 and Δ11-desaturases, because these enzymes play an important role in moth pheromone biosynthesis (Liénard *et al.* 2014; Yew & Chung 2015). Previous work with *B. anynana* suggested that a Δ11-desaturase is involved in the production of MSP1 (Liénard *et al.* 2014). Both Δ9- and Δ11-desaturase were present in the transcriptome, mainly in antennae. A phylogenetic tree containing our Δ9- and Δ11-desaturase contigs further revealed a similar position within the larger desaturase phylogenetic tree, compared to earlier work (Liénard *et al* 2014; Supplementary Figure 2).

To get more insight into the role played by the Δ9 and Δ11 desaturase gene, we used RT-qPCR (cf. protocol in Arun *et al* 2015) to compare transcript abundance between different adult wing tissues, the main tissue producing MSP1 (and using RNA extracted from new samples). Δ9-desaturase transcript abundance was approximately four-fold higher than that of Δ11-desaturase (Welch Two Sample t-test on normalized Ct values; t = −8.56, df = 9.11, p-value <0.0001; Supplementary Figure 1). When comparing the spatial distribution of MSP1 on *B. anynana* body parts with our RT-qPCR data for the two Δ-desaturase genes (Supplementary Figure 3A), the expression profile of the Δ9-desaturase gene, but not the Δ11-desaturase gene, matched MSP1 distribution (Supplementary Figure 3B and 3C, respectively). Indeed, the Δ9-desaturase gene showed overall significant variation in transcript abundance across tissues that correlated with the distribution pattern of MSP1 (nested ANOVA, n = 24 samples with 3 biological replicates and 2 technical replicates; F_3,6_ = 883.5; p-value < 0.01). Specifically, Δ9-desaturase was found to be significantly overexpressed in male wing parts containing the androconia that produce MSP1, compared to remaining male wing tissues (nested ANOVA; F_1,4_ = 1,814.0; p-value < 0.01) and female wings (nested ANOVA; F_1,4_ = 50.5; p-value < 0.01). Moreover, Δ9-desaturase gene expression was also found to be significantly overexpressed in male head tissue containing MSP1. No such match between gene expression and MSP1 abundance was found for the Δ11-desaturase gene, which showed no significant variation in transcript abundance across tissues known to contain MSP1 (nested ANOVA, F_3,6_ = 0.07; p-value < 0.01). Altogether, these findings suggest that a Δ9 desaturase plays a role in *B. anynana* pheromone biosynthesis.

We then searched for genes from a second gene family known to be involved in sex pheromone production in *B. anynana*: fatty acyl reductases (*far*), which convert fatty-acyl pheromone precursors to alcohol (Liénard *et al.* 2014). While more than 20 FARs have been experimentally characterized from 23 moth and butterfly species, all FARs implicated in moth and butterfly sex pheromone biosynthesis are restricted to a single clade, suggesting that one FAR group was exclusively recruited for pheromone biosynthesis (Tupec et al. 2017; 2019 and refs therein). In our transcriptome, two reductase contigs were annotated and identified in male and female antennae: enoyl-CoA reductase and fatty-acyl reductase 1, *far 1*. As *far1* and another fatty-acyl reductase, *far2*, were previously found to be involved in MSP2 and MSP1 biosynthesis, respectively (Liénard *et al.* 2014), we manually mined our transcriptome for *far1* and *far2* contigs by n-blasting *far1* and *far2* specific gene sequences. Contigs matching *far1* were largely overexpressed in androconia (171 copies), compared to wing controls (0 copies; Supplementary Table 2). While contigs matching *far2* showed an overall low expression level in wing tissues (Supplementary Table 3), a previous qRT-PCR study revealed that *far2* gene expression matched MSP1 biosynthesis (Arun *et al.* 2015), highlighting the potential importance of *far2* for *B. anynana* pheromone production.

The low expression level of *far2* is surprising given the amount of MSP1 present on male wings (2 μg/individual on average); hence we suggest that alternative candidates for MSP1 biosynthesis could be aldo-keto reductases, two of which are among the most expressed genes in androconial male wing tissues (Supplementary Table 1). Indeed, fatty-acyl reductases are usually associated with the reduction of aldehyde into alcohols producing various sex pheromone components in moths (Moto *et al.* 2003; Ando *et al.* 2004), but aldo-keto reductases are regularly found highly expressed in sex pheromone transcriptomes of moth species (Gu *et al.* 2013; Zhang *et al.* 2014). Guo *et al.* (2014) and Yamamoto *et al.* (2016) have proposed that aldo-keto reductases are involved in sex pheromone biosynthesis in the moths *Helicoverpa armigera* and *Bombyx mori* by reducing 9-hexadecenal, 11-hexadecenal and 10E,12Z-hexadecadienal into alcohol. Our expression data suggest that an aldo-keto reductase, with or without *far2*, may be involved in MSP1 biosynthesis.

### B. anynana *sex pheromone reception*

The genomes of the butterflies *Danaus plexippus* and *Heliconius melpomene* (i.e., species for which phylogenies of odorant receptor genes were available) have revealed a large number of genes belonging to families known to be involved in olfactory reception in moths, including odorant receptors, and odorant binding proteins (Zhan *et al.* 2011; Dasmahapatra *et al.* 2012; Briscoe *et al.* 2013). Specifically, the odorant receptor (OR) and odorant binding protein (OBP) gene families contain lineages specialized in the detection of sex pheromones in moths, the so-called pheromone receptors (PRs) and pheromone-binding proteins (PBPs) (Sakurai *et al.* 2004; Zhang & Löfstedt 2015; Bastin-Héline et al 2019; Vogt et al 2015). ORs are transmembrane receptors that bind volatile chemicals and are responsible for signal transduction in insect olfactory sensory neurons. They exhibit various response tuning breadths, and moth ORs involved in pheromone detection are often (but not always) highly specific to one or a few pheromone components (Zhang & Löfstedt 2015). We, therefore, expected to identify ORs binding to each of the three known chemical components of the *B. anynana* male sex pheromone: MSP1, 2, and 3 (Nieberding *et al.* 2008). We identified the obligatory co-receptor “Orco” and 16 ORs in the transcriptome, some of which were overexpressed in antennae compared to other adult tissues (Supplementary Table 4). Phylogenetic analysis revealed that ORs expressed in *B. anynana* were distributed among various lepidopteran OR lineages (Figure 2; Dasmahapatra *et al.* 2012), but none were located in the classically defined sex pheromone receptor clade (Bastin-Héline *et al.* 2019; Shen *et al.* 2020; Figure 2). Our results suggest that *B. anynana* sex pheromone reception may have evolved from lepidopteran OR lineages other than the sex pheromone lineage.

**Figure 2:**
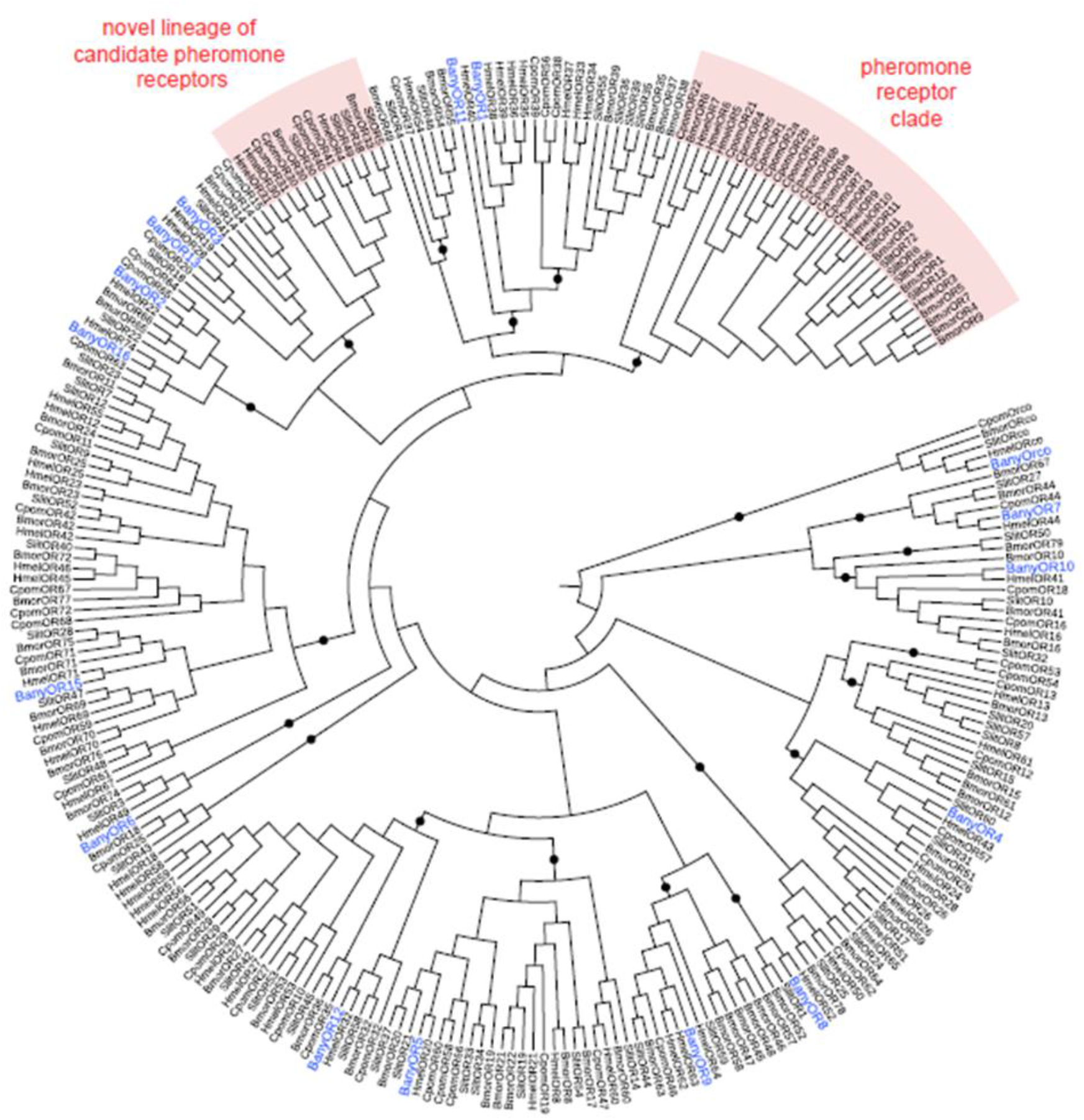
Maximum-likelihood phylogeny of Lepidopteran odorant receptors (OR), including the 16 ORs found in the B. anynana transcriptome (BanyOR, in blue). Bmor, Bombyx mori; Cpom, Cydia pomonella; Hmel, Heliconius mefpomene; Slit, Spodoptera fittorafis. Black circles indicate branchings highly supported by the approximate likelihood-ratio test (aLRT > 0.95).

Recent studies have revealed that moth PRs do not constitute a monophyletic clade and, instead, evolved several times during OR evolution (Yuvaraj et al 2017, Bastin-Héline et al 2019). Functional PRs that have been found outside of the PR clade in some moth species were identified based on their sex-biased expression. We, therefore, searched for potential *B. anynana* PRs by quantifying the expression levels between sexes using qPCR, expecting that PR in *B. anynana* should be more expressed in male than female antennae. We further expected that gene expression levels would correlate with temporally varying physiological and biological needs. In moth species, PRs are critical for detecting the female sex pheromone and the male’s behavioral and physiological responses to female sex pheromones were shown to be affected by moth age and mating status (Soques *et al.* 2010, Saveer et al 2012). We, therefore, tried to identify *B. anynana* candidate PRs by comparing RNA expression levels in females with different mating status (using qPCR). We expected that virgin females that had developed either in isolation (naive “virgin”) or in the presence of male scent (“virgin sensitized”) would exhibit higher expression levels for OR genes responsible for detecting the male sex pheromone, compared to mated females (“mated”)(Zeng et al 2013 Biochem Biophys Res Commun; Immonen et al 2012 PRSB). This difference would be due to virgin females taking information about the composition of the male sex pheromone for choosing mates regarding their inbreeding level (van Bergen *et al.* 2013) or their age (Nieberding *et al.* 2012), and because recently mated females are much less receptive to courtship attempts in *B. anynana*. The candidate genes Ban_OR1, Ban_OR2 and Ban_Orco were selected for qPCR experiments because these genes displayed the highest expression among the 16 identified candidate ORs and were significantly overexpressed in antennae compared to control libraries (Supplementary Table 4). Orco expression was significantly decreased in mated compared to virgin (naïve or sensitized) females, but Bany_OR1 or Bany_OR2 were not (Figure 3). This finding suggests that the regulation of the expression of Orco could be a mediator of sex pheromone receptivity. Orco, and not specific parts of the odorant receptor dimer, such as OR1, OR2 or other odorant receptors that we did not test here, could be regulated by sex pheromone communication, similar to what was previously found in cockroaches (Soques *et al.* 2010; Latorre-Estivalis *et al.* 2015).

**Figure 3:**
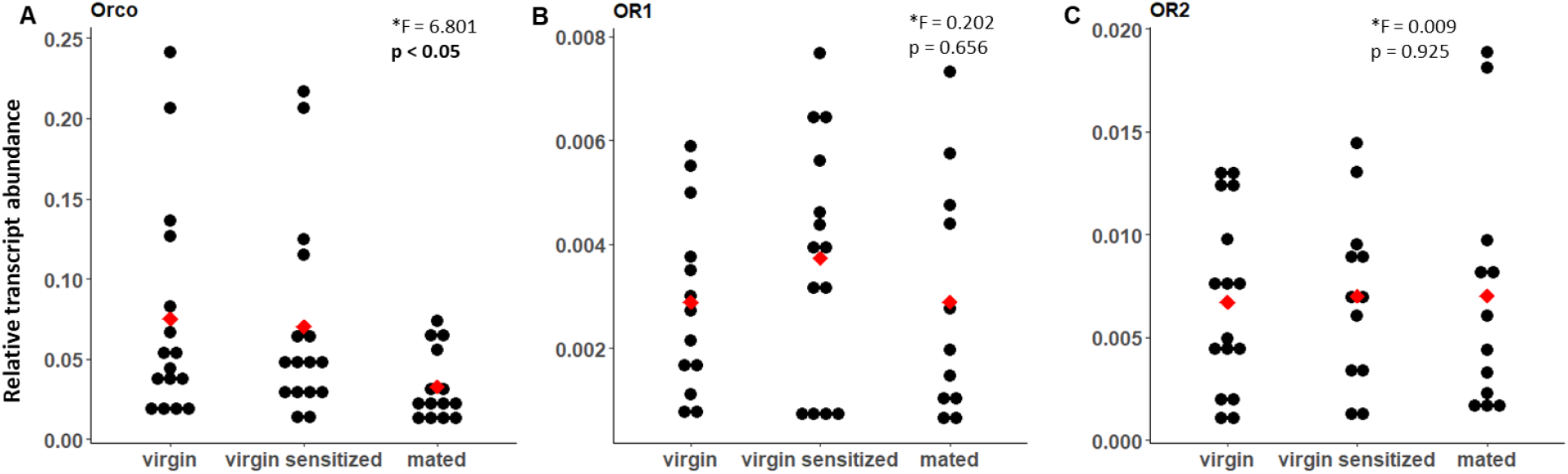
qPCR expression level of odorant receptors (OR) in antennae of female B. anynana with different mating status. Oreo (A), but not OR1 (B) or OR2 (C), RNA expression level differed significantly in virgin naïve (left) and virgin sensitized (middle) compared to mated (right) females. Each treatment is the mean of 3 to 7 biological replicates. A nested ANOVA was used to test for differences between groups. F and p values are included for each graph. *log transformed data.

In addition to the work described above, we tried to functionally investigate if some specific OR candidate genes were responsible for the detection of male pheromone components using heterologous expression in *D. melanogaster* olfactory sensory neurons coupled to electrophysiological recordings. These experiments did not lead to functional validation, but the procedures followed and results obtained are described in Supplementary File 1.

A second gene family specific to insects, the OBP family, is involved in olfaction by solubilizing semiochemicals once they have entered the aqueous lymph within olfactory sensilla (Leal et al 2013). OBPs have been proposed to play an important role in response sensitivity. In Lepidoptera, a dedicated lineage of OBPs (the so-called “pheromone-binding proteins” or PBPs) has evolved high affinity towards pheromone components (Vogt *et al.* 2015). We identified 46 contigs assembled into 13 OBP unigenes expressed in our *B. anynana* transcriptome (Supplementary Table 5), a number lower than what has been described in various transcriptomes from moth species (49 predicted OBPs in *Spodoptera littoralis* and *Manduca sexta;* Walker et al 2019) and in the genomes of two butterfly species (32 in *Danaus plexippus*, 51 in *Heliconius Melpomene*; Pelosi *et al.* 2014, 2018; Venthur & Zhou 2018). *B. anynana* expressed OBPs were found in most subclades of the phylogenetic tree of lepidopteran OBPs, including general odorant binding protein 1 and 2 lineages, as well as classic, minus-C, plus-C and duplex OBP lineages (with categories based on the level of sequence homology and conserved amino acid signatures; Supplementary Figure 4). In Lepidoptera, the OBP gene family also includes a lineage of the PBPs, thought to transport pheromone molecules (Vogt *et al.* 2015). In moths, such as *Manduca sexta* and *Bombyx mori*, trichoid sensilla are associated with pheromone perception and express specifically PBP-A. No *B. anynana* expressed OBP clustered in the pheromone-binding protein −A or −B lineages (Supplementary Figure 4). This is similar to what has been found in other butterfly species: the PBP-A lineage is lacking in the genome of *Danaus plexippus* and the PBP-A and PBP-B lineages are absent from the genomes of *Heliconius melpomene* and *Melitaea cinxia* (Vogt *et al.* 2015). In contrast, we did find two candidate PBPs (Supplementary Table 5), expressed in *B. anynana* antennae, which belong to the pheromone-binding protein-C and −D lineages present in all butterfly genomes investigated to date (Vogt *et al.* 2015). These candidate PBPs indeed correspond to the two sole candidate PBP genes identified in the *B. anynana* genome, and are both most similar to two PBPs found in the antennae of *H. melpomene* (Dasmahapatra et al. 2012; Supplementary Figure 4). In most moths, PBP-C and PBP-D OBPs are expressed in basiconic sensilla and are associated with foraging (Vogt *et al.* 2015). Although we cannot exclude that we missed BanOBPs in our transcriptome, the lack of a PBP-A subgene family in *B. anynana*, as in the four other butterflies studied to date (*H. melpomene, D. plexippus, M. cinxia, P. polytes*), suggests that butterflies lost this gene lineage (at least in the Nymphalidae, the family of butterflies to which the sampled species belong), and possibly use other PBP lineages to functionally aid the OR-pheromone connection. The transcriptome was further mined for Chemosensory Proteins (CSPs), a third gene family potentially implicated in olfaction in insects (Sánchez-Gracia *et al.* 2009; Vieira & Rozas 2011), for which the results are available in Supplementary Table 6.

### B. anynana *sex pheromone regulation*

Eleven contigs were found overexpressed in male compared to female brains, but their role in the regulation of sex pheromone processing remains open (Supplementary Table 1). Given its role as a key regulator of female sex pheromone biosynthesis in many moth species (Bloch *et al.* 2013), we focused our attention on Pheromone Biosynthesis Activating Neuropeptide (PBAN). We hypothesized that PBAN could be involved in male sex pheromone regulation in *B. anynana*, and looked for it in our transcriptome database. We identified one unigene annotated as PBAN (BA_PBAN.1.1), which was expressed in adult heads. We used this sequence to obtain the complete cDNA sequence of PBAN in *B. anynana* (RACE), Ban_PBAN (Figure 4A). The phylogenetic reconstruction of PBAN across Lepidoptera shows monophyly of butterfly PBANs (data not shown). We next investigated the PBAN cDNA tissue distribution using semi-quantitative and quantitative PCR. PBAN was found to be expressed in adult heads, but not in other tissues, and more strongly in males than in females (Figure 4B). PBAN in male moths is suspected to be involved in male pheromone biosynthesis: the PBAN receptor of the moth *Helicoverpa armigera* was found expressed in male hairpencils, and PBAN stimulation of the hairpencils was found to be responsible for the production and release of male pheromonal components (Bober & Rafaeli 2010). Next, we used RT-qPCR and found that PBAN expression level in male brains correlated with the amount of male sex pheromone found on male wings during the adult male’s lifetime, with maximum content around 15 days (Figure 4C; Nieberding *et al.* 2012).

**Figure 4:**
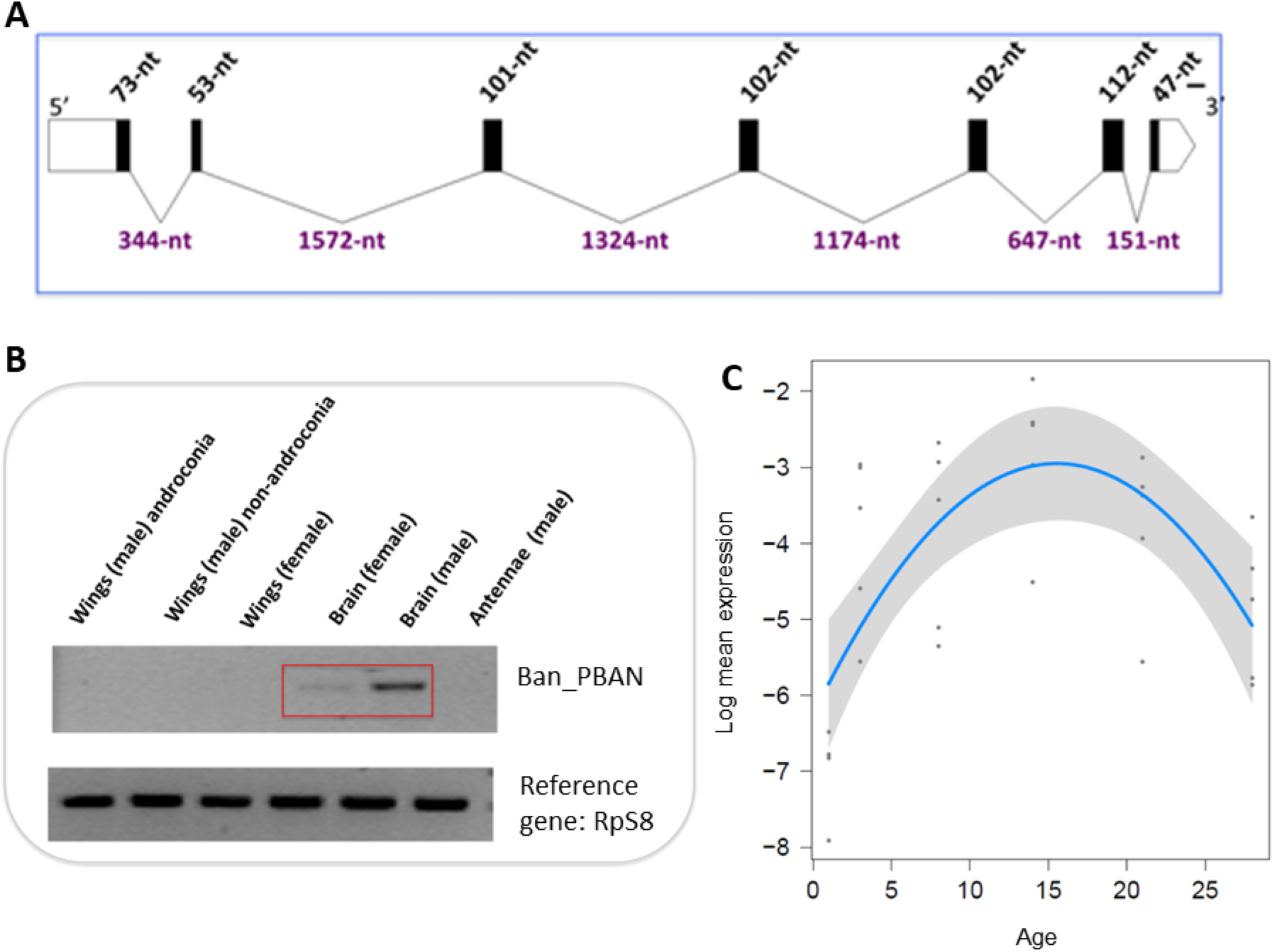
Pheromone binding activating neuropeptide (PBAN) expression in 8. anynana. A) Structure of Ban_PBAN full length gene sequence (594 bp). The 7 exons (encoding 198 amino acids) are represented by black boxes and the 6 introns by lines. B) PBAN expression level quantified by semi-quantitative PCR in adult tissues (brains, wings, antennae) of males and females ranging from 3 to 14 days of age. Higher levels of PBAN are observed in male brains compared to the other adult tissues. C) PBAN expression level quantified by real time qPCR in adult male brains from 1 to 28 days of age.

In moths, production of volatile sex pheromones usually shows a circadian pattern that is regulated by PBAN and correlates with the female “calling” behavior (extrusion of the sex pheromone gland) during specific hours of the scotophase (Rosén 2002; Bloch, Hazan, and Rafaeli 2013). A circadian rhythm of male sex pheromone production was also found in the moth *Aphomia sabella* (Levi-Zada *et al.* 2014). We tested whether *B. anynana* displayed daily variation in courtship activity, MSP production, and PBAN expression in 8-day old adult males. We found that courtship activity peaked 7 to 12 hours after sunrise, and courtship activity was significantly higher in the afternoon compared to the rest of the day (Figure 5A; likelihood ratio test on the “Time²” effect, p= 0.005; Supplementary Table 7 and 8). Similarly, MSP production significantly varied during the course of the day, and peaked around maximum courtship activity, with MSP1/MSP2 and MSP2/MSP3 ratios displaying significant reversed changes during the day (Figure 5D; likelihood ratio test on the “Time² “p= 0.004 and p = 0.002, respectively; Supplementary Table 8). MSP amounts also displayed a slight, but non-significant, variation with time of the day (Figure 5C; likelihood ratio test on the “Time²” effect, 0.007 < p < 0.09; Supplementary Table 8). MSP titers were estimated to be minimum around 11 hours after sunrise for MSP1 and MSP3, while the MSP2 titer was estimated to be at a maximum 12.4 hours after sunrise. We further found that PBAN expression significantly varied throughout the day as well (Figure 5B), with the highest expression 11 to 14 hours after sunrise. Daily variation in PBAN expression thus correlates both to male courtship activity and to male sex pheromone quantities found on male wings: all three traits peak during the afternoon and PBAN expression is maximal just before the peak in MSP2/MSP1 and MSP2/MSP3 ratios and MSP2 amount. This suggests that the daily regulation of male sex pheromone may be associated to circadian variation in PBAN expression, a neuropeptide that is specific to sex pheromone regulation in moths (Cheng *et al.* 2010).

**Figure 5:**
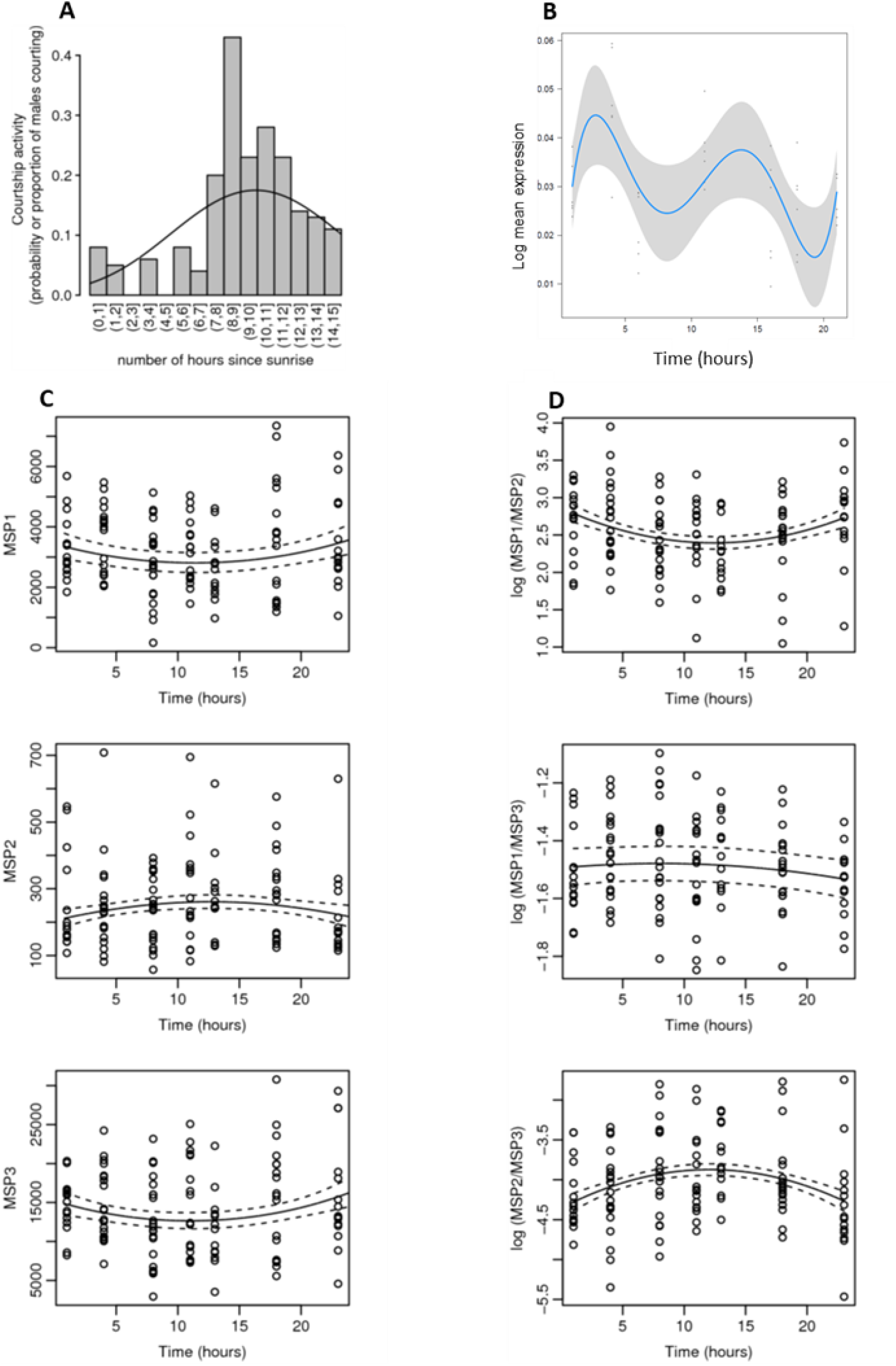
Daily variation in *B. anynana* male courtship activity (A), PBAN expression by qPCR (B), MSP production (C) and log MSP ratio production (D).

In addition to the work described above, we aimed to functionally demonstrate the role of PBAN expression in regulating male sex pheromone biosynthesis in *B. anynana*. These experiments did not lead to functional validation of the role of PBAN, but all procedures followed and results obtained are described in Supplementary File 2.

### Conclusions and perspectives

This is the first time a transcriptomics approach has been used to gain insight into the different aspects of sex pheromone communication in a butterfly. Our model, *B. anynana*, is the only butterfly species for which the sex pheromone composition and its role in male mating success in relation to different fitness-associated traits (e.g., age and inbreeding level) have been demonstrated (Costanzo & Monteiro 2007; Nieberding *et al.* 2008, 2012; van Bergen *et al.* 2013). The male sex pheromone composition and function in communication between sexes has been partially analyzed in only a few additional butterfly species, including *Pieris napi*, *Colias eurytheme*, *Danaus gilippus*, *Idea leuconoe*, and more recently *Heliconius melpomene* (Pliski & Eisner 1969; Taylor 1973; Grula *et al.* 1980; Sappington & Taylor 1990a, b; Nishida *et al.* 1996; Andersson *et al.* 2007; Darragh *et al.* 2017).

Mosaic evolution appears to have taken place at the molecular level based on our investigation of the pathways involved in the production, reception, and regulation of the sex pheromone in *B. anynana*: our data suggest that the biosynthesis of the three chemical components forming the male sex pheromone could be partly due to moth-specific genes (*far1* and *far2* for the MSP2 and MSP1 components, respectively) and partly due to genes present in insects other than moths (Δ9-desaturase, aldo-keto reductase for the MSP1 component). This is also likely the case for the MSP3 component whose synthesis is not expected to rely on moth-specific gene families, as this pheromone component is not derived from fatty acids. None of the expressed odorant receptors or odorant binding proteins in *B. anynana* belonged to Lepidoptera-specific gene lineages responsible for sex pheromone reception in moths, suggesting that sex pheromone reception in this butterfly may have evolved independently from their moth ancestors. In contrast, we found that sex pheromone biosynthesis could be regulated by the neuropeptide PBAN in both moths and butterflies, an evolutionarily shared derived trait for Lepidoptera.

Recently, the genomes of 250 species of skippers (Hesperiidae; Li et al. 2019) and 845 North American butterfly species (Zhang *et al.* 2019) have been sequenced. A systematic comparative analysis of major gene families involved in moth sex pheromone communication in these ~1000 butterfly genomes would provide important information on the level of conservation of molecular pathways when butterflies diverged from moths about 98 million years ago.

## Materials and methods

### Insects

*Bicyclus anynana* (Butler, 1879) (Lepidoptera: Nymphalidae) originated from an outbred wild type population that was collected in Malawi, Africa, in 1988 using 80 gravid females. Since 1988, several hundred individuals have been used each generation to maintain high levels of heterozygosity (Brakefield *et al.* 2001) in a climate-controlled room at a temperature of 27ºC, a relative humidity of 70%, and a photoperiod of L:D 12:12. Larvae were kept under these conditions on maize plants *Zea mays*, and adults fed mashed banana *Musa acuminate* for all experiments, except when stated otherwise.

### Transcriptome

#### Tissue collection

For the transcriptome dataset, several hundreds of virgin males and females were separated at the pupal stage in different cages, and tissues collected in March 2010. Pupal tissues were collected from male and female pupae 1 to 7 days after pupation (1 or 2 pupae per day after pupation and per sex); the wing imaginal discs were dissected as described in (Brakefield *et al.* 2009). Tissues for adult libraries (wings, heads and antennae) were collected from adult virgin males and females aged 1, 3, 5, 8, 10 and 14 days after emergence (Supplementary Figure 5): ~50 adults were used per age category and per library for wing libraries, ~10 adults were used per age category and per library for head tissues, and ~5 adult females and 5 males were used per age category for the antennae library. Brain tissue was obtained by cutting the head and cutting off the eyes, the proboscis and the antennae; antennal tissue was collected for a similar number of adult males and females (Figure 1). Dissected tissues were conserved immediately at −20°C in RNAlater (Sigma-Aldrich).

#### RNA extraction

RNAs of all dissections were extracted in April 2010, within a month after collection of tissues, in an RNA-free environment, on ice, and using the RNeasy Mini kit and the RNAase free DNAase kit (Qiagen). After RNA extraction, 1 μl of each RNA extract was used for testing RNA quality and quantity using a Bioanalyzer Systems (Agilent) at the LUMC hospital in Leiden (the Netherlands, courtesy of Dr Jeroen Pijpe), and using the RiboGreen RNA quantification kit, respectively. The remaining RNA extract was stored at −80°C for cDNA synthesis. For cDNA synthesis, we first pooled all RNA extracts dedicated to the same library in one tube per library, in such a way that: *i*) the same amount of RNA was present for each sex (male and female), *ii*) each life stage was represented by similar RNA amounts (days 1 to 7 after pupation for pupal tissue libraries, days 1 to 14 after emergence for adult tissue libraries; Supplementary Table 9). Total RNA yield was 27 to 40 μg per library as requested for sequencing.

#### mRNA isolation, cDNA synthesis and sequencing

mRNA capture, cDNA synthesis and tagging for Titanium 454-sequencing was performed by Biogenomics, a KU Leuven Research & Development Division of the Laboratory of Animal Diversity and Systematics (Leuven, Belgium). Between 370 and 1,340 ng (0.3 and 1.6%) mRNA yield was obtained for each library, providing enough mRNA (minimum 200 ng per library) for cDNA construction and tagging. Yet, cDNA synthesis failed when started from mRNA, which is why a SMART cDNA synthesis was performed from total RNA. A custom normalization step (based on the EVROGEN Trimmer kit) was optimized in collaboration with the Roche R&D department and applied to the cDNA libraries, as no validated normalization protocol was available from Roche in 2010 for Titanium cDNA sequencing. Each normalized library was quality checked for fragment length and integrity before sequencing. Each library was subjected to GS FLX Titanium Emulsion PCR and Sequencing, and each library was sequenced 5 times. After sequencing, data were processed through certified Roche software (GS Transcriptome Assembler/Mapper) and custom scripts for advanced analysis. Basic data analysis included read quality trimming and assembly into contigs, including potential alternative splicing products. The sequences were trimmed by removing low quality sequences, ambiguous nucleotides, adapter sequences, and sequences with lengths less than 20 nucleotides. The 454-sequencing generated 824,439 reads, with an average length of 293 base pairs and a total of 242,005,027 nucleotides (Supplementary Figure 6).

#### Transcriptome assembly, quantification, and annotation

Adaptors were removed with smartkitCleaner and adaptorCleaner. Raw sequences (reads) were cleaned with the software Pyrocleaner (Mariette et al. 2011[2]), using the following criteria: (1) complexity/length ratio less than 40 (using a sliding window approach based on 100 bp sequence length, and a step of 5 bp); (2) duplicate read removal (see bias associated with pyrosequencing, due to the random generation of duplicate reads); (3) removal of too long/too short reads (maximum and minimum read length = mean read length ± 2 SD); (4) removal of reads with too many undetermined bases (more than 4 %). Contaminations were discarded by searching hits against *E. coli*, phage and yeasts.

The reads were assembled *de novo* in 43,149 contigs of 488 base pairs on average with a total of 21,087,824 nucleotides (Supplementary Figure 6). The average GC content was 36.44%. The assembly was perform with tgic l (https://academic.oup.com/bioinformatics/article/19/5/651/239299) version 2.1 using standard parameters. The reads where realigned to the contigs and singletons with bwa aln version 0.7.2 using standard parameters and transformed in bam format, sorted and indexed with samtools version 0.1.19 with default parameters The bam files were then processed with samtools idxstats to extract expression measures in the form of numbers of reads aligned on each contig for every condition. These measures were than merged to produce the quantification file using unix cut and paste commands. Diamond was used to search for sequence homology between contig and the following generalist databases: UniProtKB/Swiss-Prot, UniProtKB/TrEMBLRelease of April, NR release of end of March.

The following species from the ensemble database were queried: *Bicyclus anynana* (nBA.0.1), *Calycopis cecrops* (v1.1), *Danaus plexippus* (v3), *Heliconius melpomene melpomene* (Hmel2.5), *Junonia coenia* (JC v1.0), *Lerema accius* (v1.1), *Melitaea cinxia*, *Papilio machaon* (Pap_ma_1.0), *Phoebis sennae* (v1.1) and *Pieris napi* (v1.1).

#### Identification of specific gene families

We also mined the transcriptome for specific families of genes supposedly involved in sex pheromone communication based on the available evidence in moths (Lepidoptera): desaturases, reductases, odorant receptors (OR), odorant binding proteins (OBP), and chemosensory proteins (CSP). For this:

i. we downloaded the DNA sequence of every *B. anynana* contig named as a desaturase, reductase, OR, OBP or CSP in our transcriptome;
ii. we checked the homology of the sequence of each candidate contig with gene members of the same family in other Lepidoptera by performing a blastx in Genbank;
iii. every *B. anynana* contig that showed significant homology in step *ii* was blasted in the transcriptome, allowing us to find more *B. anynana* ESTs of the same gene family, even if some had not been annotated as such. All these contigs and ESTs were then “candidate members of each respective gene family”. If no significant homology was found using blastx in step *ii*, the sequence was removed from the list of candidate members of the gene family;
iv. every *B. anynana* contig and EST candidate was then translated into an amino acid sequence using Expasy (https://web.expasy.org/translate/). When necessary, cdd analyses of domains were done. Using this procedure, 27 OR, 44 OBP and 70 CSP candidate members were found in the *B. anynana* transcriptome (Supplementary Tables 4, 7 and 8 for OR, OBP and CSP, respectively; for reductases and desaturases see Results). For example, 40 contigs were initially annotated as “odorant-binding protein” in our transcriptome, based on the characteristic hallmarks of the OBP protein families, including six highly conserved cysteines, i.e., “C” (in Lepidoptera C1-X25-30-C2-X3-C3-X36-42-C4-X8-14-C5-X8-C6, with “X” being any amino acid; Xu *et al.* 2009). As sequence conservation between OBPs is low, i.e., between 25 and 50% identity for amino acid sequences, manually mining the transcriptome allowed us to find another seven OBP candidate members (Supplementary Tables 4, 7, and 8 for OR, OBP and CSP, respectively).
v. Candidate members were then manually aligned in Bioedit to group them into distinct expressed gene units, or unigenes: 17 Bany_OR unigenes, 9 Bany_OBP unigenes, including in some cases more "gene subunits" when contigs were similar enough to suggest that they represented different allelic variants of the same gene, such as Bany_OBP3, Bany_OBP4, Bany_OBP6 (Supplementary Table 5) and 8 Bany_CSP unigenes with some more gene subunits as well (Supplementary Table 6).
vi. The expression level of each candidate unigene across libraries was then obtained by pooling the number of copies in the *B. anynana* transcriptome of each EST and contig forming the unigene.

### Phylogenies

For the OR phylogeny, amino acid sequences found in the *B. anynana* transcriptome were aligned with OR sequences previously identified in the genomes of *B. mori* and *H. melpomene* (Dasmahapatra *et al.* 2012) and in antennal transcriptomes of *C. pomonella* (Walker *et al.* 2016) and *S. littoralis* (Walker *et al.* 2019). Alignment was performed with MAFFT v7 (https://mafft.cbrc.jp/alignment/server/), and the maximum-likelihood phylogeny was built using PhyML 3.0 (Guindon *et al.* 2010). Branch support was assessed using a likelihood-ratio test (Anisimova & Gascuel 2006). Published datasets of Lepidoptera protein sequences from previous phylogenetic studies were used for testing the phylogenetic position of *B. anynana* ORs (Poivet *et al.* 2013; Supplementary File 3), OBPs (Vogt *et al.* 2015; Supplementary File 4), desaturases and reductases (Liénard *et al.* 2014; Supplementary Files 5 and 6).

### Quantitative real time PCR

Biological replicates were the mRNA extracted from either a single individual or formed by pooling 3 to 5 individuals of various ages in experiments for the “reception” and the “production” communication steps, respectively. Each treatment is represented by 3 to 7 biological replicates. The protocol used for quantitative real time PCR is described in Arun *et al.* (2015). Briefly, total RNA was extracted using the RNeasy Mini kit following manufacturer’s instructions. Residual DNA was removed after treating extracted RNA using a DNase enzyme. A nanodrop ND-1000 spectrophotometer was then used to assess the integrity of the RNA before conversion into cDNA. qRT-PCR was carried out using the SYBR green dye in a 96-well thermocycler with parameters described in Arun *et al.* (2015). Primer sequences for all genes are available in Supplementary Table 10. Relative transcript abundance was calculated using the 2(-ΔCt) method. Statistical significance of differences in expression levels expressed in Rq values after log-transformation was tested using nested ANOVA with technical replicates nested with biological replicates; the model was log (Rq) ~ treatment / biological replicate / technical replicate + Error (tissue / biological replicate / technical replicate). R version 3.6.1 (R Core Team 2020) was used for statistical analyses.

### Behavioral experiments

#### Mating experiments for quantifying odorant receptor expression levels

Naïve virgin females were reared in isolated conditions (devoid of the male secondary sexual traits putatively involved in sexual communication, i.e., olfaction, vision and audition) directly after egg collection. The virgin sensitized females were reared in a MSP-containing environment near cages containing males. The sensitized mated females were reared in a MSP-containing environment and mated at an age of 3 days. All females were sacrificed at day 5 and the antennal tissues were used for RNA extraction and RT-qPCR analysis (as described above).

#### Daily variation in courtship activity

We tested whether courtship activity in *B. anynana* males varies throughout the day. A large number of individuals were reared and age after emergence was recorded. The day before the experiment, five males and four females between 2 and 12 days old were randomly chosen and grouped in a cage (40cm x 30cm, cylindrical). The cages were placed in a room with a temperature of ~27°C with natural light, and a 14:10 day-night regime. The butterflies were fed with banana slices and had access to water during the course of the experiment. We used five cages per trial and produced three trials with different individuals. A generalized mixed model with binomial error distribution was used to characterize the courting activity of males during the day. The presence/absence of courtship behavior for each male during 15 minutes of observations per hour was used as the dependent variable. As we expected courtship activity to peak at some time point during the day, we used “time of the day” (in the number of hours after natural sunrise) and its second order polynomial as a fixed explanatory variable. The age of males (in days) was also included as a fixed cofactor to control for the effect of age. The identity of each individual, cage and trial were used as random effects with individual nested within cage and cage nested within trial. We tested the model parameters with type III likelihood ratio tests, in which a model without the explanatory variable of interest is compared to the full model, both models being estimated by Maximum Likelihood.

#### Daily variation in male sex pheromone production

A number of butterfly couples were set up using adult virgin stock males and females. Three families were started from three couples that produced over 200 offspring. The three families were each partly reared into two different climate rooms that differed in the onset of artificial daylight (one at 9:00 am, the other at 12:00 pm). This allowed us to control for the potential effect climate cell-specific conditions on variation in MSP production. Forty to 80 males that emerged the same day were selected per family. MSP production of 8-day old males was sampled, an age at which each MSP component is produced in measurable quantities (Nieberding *et al.* 2008). Four to 7 males of each family were killed and conserved at −80° for subsequent pheromone analysis at seven sampling points during the day: 1, 4, 8, 11, 13, 18 and 23 hours after the onset of daylight. MSP production was measured as described below in the section “male sex pheromone quantification”. We used mixed models with normal error distribution to characterize the variation of MSP production during the day. The titre of each MSP component and the ratios between pairs of MSP components were used as dependent variables. MSP titres were square root transformed and MSP ratios were log-transformed to improve the normality and homoscedasticity of the residuals. As we suspected MSP production to peak at some time point during the day, we used a second order polynomial equation with time and time² as a fixed explanatory variables and family as a random effect. We tested model parameters with type III likelihood ratio tests, in which a model without the explanatory variable of interest is compared to the full model, both models being estimated by Maximum Likelihood. We estimated the percentage of variation explained by the models and each of their components with pseudo R² based on ratios of sums of squared residuals. We followed Legendre & Legendre (1998) for the variance decomposition procedure.

### Quantification of male sex pheromone production

MSP concentrations were determined as previously described (Nieberding *et al.* 2008, 2018). In short, one forewing and hindwing of each male was soaked in 600ul of hexane during 5 minutes. One ng/ul of the internal standard (palmitic acid) was then added. Extracts were analyzed on a Hewlett-Packard 6890 series II gas chromatograph (GC) equipped with a flame-ionization detector and interfaced with a HP-6890 series integrator, using nitrogen as carrier gas. The injector temperature was set at 240 °C and the detector temperature at 250 °C. A HP-1 column was used and temperature increased from the initial temperature of 50 °C by 15 °C/min up to a final temperature of 295 °C, which was maintained for 6 min.

## Acknowledgments

CMN’s team was supported by the Fonds National de la Recherche Scientifique (FNRS), FRFC grants 2.4600.10 and 2.4560.11 and CR grant 29109376, as well as UCLouvain ARC grant n◦ 10/15-031 and FSR grant n◦372 605031. BV was supported by FNRS CR grant 24905063. EJJ, NM and LBH were funded by the French National Research Agency (ANR-16-CE02-0003-01 and ANR-16-CE21-0002-01 grants).

## Acknowledgments

We would like to thank Prof. Ken-Ichi Moto and Prof. Tetsu Ando for discussions about sex pheromone biosynthesis in Lepidoptera; Dr Jeroen Pijpe for mentoring us about, and providing access to, a Bioanalyzer Systems (Agilent) at the LUMC hospital in Leiden (the Netherlands); Marleen van Eijk for assistance with preparing the butterfly tissues. This is publication BRC321 of the Biodiversity Research Center.

## Abbreviations

MSP: male sex pheromone
OR: odorant receptor
OBP: odorant binding protein
PBAN: pheromone biosynthesis activating neuropeptide

